# Butyrate differentiates permissiveness to *Clostridioides difficile* infection and influences growth of diverse *C. difficile* isolates

**DOI:** 10.1101/2022.05.20.492898

**Authors:** Daniel A. Pensinger, Andrea T. Fisher, Horia A. Dobrila, William Van Treuren, Jackson O. Gardner, Steven K. Higginbottom, Matthew M. Carter, Benjamin Schumann, Carolyn R. Bertozzi, Victoria Anikst, Cody Martin, Elizabeth V. Robilotti, JoMay Chow, Rachael H. Buck, Lucy S. Tompkins, Justin L. Sonnenburg, Andrew J. Hryckowian

## Abstract

A disrupted “dysbiotic” gut microbiome engenders susceptibility to the diarrheal pathogen *Clostridioides difficile* by impacting the metabolic milieu of the gut. Diet, in particular the microbiota accessible carbohydrates (MACs) found in dietary fiber, is one of the most powerful ways to affect the composition and metabolic output of the gut microbiome. As such, diet is a powerful tool for understanding the biology of *C. difficile* and for developing alternative approaches for coping with this pathogen. One prominent class of metabolites produced by the gut microbiome are short chain fatty acids (SCFAs), the major metabolic end products of MAC metabolism. SCFAs are known decrease the fitness of *C. difficile* in vitro and that high intestinal SCFA concentrations are associated with reduced fitness of *C. difficile* in animal models of *C. difficile* infection (CDI). Here, we use controlled dietary conditions (8 diets that differ only by MAC composition) to show that *C. difficile* fitness is most consistently impacted by butyrate, rather than the other two prominent SCFAs (acetate and propionate), during murine model CDI. We similarly show that butyrate concentrations are lower in fecal samples from humans with CDI relative to healthy controls. Finally, we demonstrate that butyrate impacts growth in diverse *C. difficile* isolates. These findings provide a foundation for future work which will dissect how butyrate directly impacts *C. difficile* fitness and will lead to the development of diverse approaches distinct from antibiotics or fecal transplant, such as dietary interventions, for mitigating CDI in at-risk human populations.

**IMPORTANCE:** *Clostridioides difficile* is a leading cause of infectious diarrhea in humans and it imposes a tremendous burden on the healthcare system. Current treatments for *C. difficile* infection (CDI) include antibiotics and fecal microbiota transplant, which contribute to recurrent CDIs and face major regulatory hurdles, respectively. Therefore, there is an ongoing need to develop new ways to cope with CDI. Notably, a disrupted “dysbiotic” gut microbiota is the primary risk factor for CDI but we incompletely understand how a healthy microbiota resists CDI. Here, we show that a specific molecule produced by the gut microbiota, butyrate, is negatively associated with *C. difficile* burdens in humans and in a mouse model of CDI and that butyrate impedes the growth of diverse *C. difficile* strains in pure culture. These findings help to build a foundation for designing alternative, possibly diet-based, strategies for mitigating CDI in humans.

## INTRODUCTION

*Clostridioides difficile* is an opportunistic diarrheal pathogen and is an “urgent threat” to global health, as it causes over 220,000 cases and 13,000 deaths per year in the United States alone [1]. A disrupted (dysbiotic) gut microbiome, most commonly resulting from antibiotic use, is the primary risk factor for *C. difficile* infection (CDI) [2], highlighting the gut microbiome as a key mediator of CDI. Therefore, measures to positively impact the composition and function of the gut microbiome represent potential approaches to understand and mitigate *C. difficile* pathogenesis.

Diet is one of the most powerful ways to impact the composition and function of the gut microbiome [3,4]. A growing body of literature demonstrates that dietary changes impact *C. difficile*, the microbiome, and the host during animal models of CDI. For example, low protein diets are protective against CDI and high fat/high protein diets exacerbate CDI [5,6] and availability of the amino acid proline in particular impacts *C. difficile* fitness in murine models [7]. Diets containing inulin, xanthan gum, and complex mixtures of microbiota accessible carbohydrates (MACs) reduce *C. difficile* burdens below detection in mice [8,9] and fructooligosaccharides (FOS) increase survival in infected hamsters [10]. Another carbohydrate, trehalose, increases CDI mortality in mice [11] but does not impact *C. difficile* burdens or virulence in chemostats containing human-derived microbiomes [12], together highlighting the need to understand how diet influences both host- and microbiome-driven factors that impact CDI outcomes. Finally, the abundance of metals such as zinc also correlate with several measures of CDI severity in mice [13]. Together, these studies support that microbiome- and host-dependent metabolite availability in the gut, rather than a specific “susceptible” or “resistant” microbiome configuration, defines colonization resistance against *C. difficile* [8,14–16]. Furthermore, each of the aforementioned diet-driven impacts on CDI represents an opportunity to understand the diverse metabolic requirements of, and niches occupied by, *C. difficile* during CDI and are likely to lead to the development of new concepts and approaches for mitigating CDI in at-risk human populations. Notably, this previous work was carried out under controlled experimental conditions which were designed to specifically manipulate conditions of interest using animal models of CDI and a limited number of *C. difficile* strains. Therefore, though animal models of CDI recapitulate many relevant aspects of human disease, it is unclear the extent to which these findings translate to human populations who are infected by phylogenetically diverse *C. difficile* strains and who differ in important parameters like immune status and dietary habits.

Of the dietary inputs described above which impact CDI, MACs represent a particularly high-yield avenue for diet-focused work on *C. difficile*. In particular, the short chain fatty acids (SCFAs), which are the metabolic end products of MAC metabolism by the microbiome [17], impact *C. difficile* fitness in pure culture and in animal models of infection [8,18,19] and have pleiotropic beneficial effects on the host [20–26]. Three SCFAs (acetate, propionate, and butyrate) are the most abundant metabolites in the gut, together reach concentrations of over 100mM in the gastrointestinal tracts of humans [27], and are influenced by host MAC consumption. The dysbiotic conditions which favor CDI are characterized by low SCFA concentrations in both humans and animal models [8,11,15,28,29].

Despite the emerging understanding of the impact of dietary MACs and their metabolic end-products on CDI and the promise for rapid translation to humans, key questions remain. For example, which MACs are most effective in impacting CDI? What parameters differentiate effective MACs from ineffective MACs? What mechanism(s) underly these differences? Are these conclusions generalizable to all *C. difficile* strains? To begin to answer these questions, this study leverages a murine model of CDI, human samples, and a collection of *C. difficile* isolates to demonstrate that elevated concentrations of butyrate are associated with a reduction in *C. difficile* fitness in pure culture, in mice, and in humans. Together, these findings provide the foundation for future work aimed at understanding the metabolic interactions that dictate *C. difficile* fitness and pathogenesis and for developing new approaches to mitigate CDI in at-risk human populations.

## RESULTS

### Inulin and FOS differentially impact C. difficile burdens in mice

In previous work, we demonstrated that inulin, a β-2,1-linked fructan, suppresses *C. difficile* burdens in a murine model of CDI [8]. To begin to test the generalizability of these findings to other purified MAC sources, we focused on FOS, which is structurally identical to inulin except for its degree of polymerization (DP) (FOS DP = 2-8 and inulin DP = 2-60) [30]. In contrast to mice fed inulin, mice fed FOS retain high burdens of *C. difficile* 630 during CDI (**Figure 1**). These results generated two possible hypotheses, that the effect of MAC sources on *C. difficile* burden is driven by either MAC effects on the microbiota or by direct effects on *C. difficile*.

**Figure 1.**
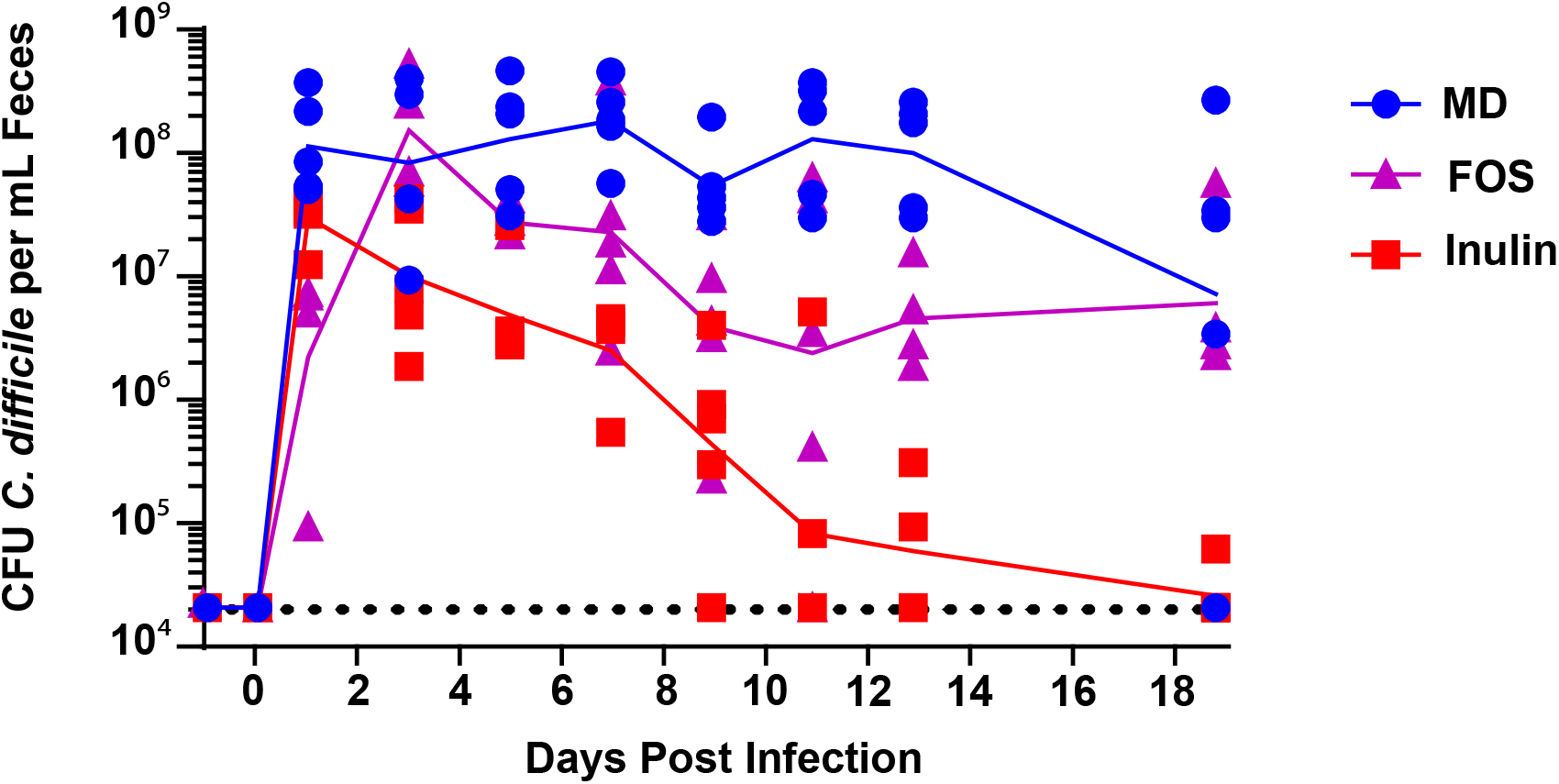
A diet containing inulin, but not FOS, as the sole MAC source reduces *C. difficile* 630 colonization below detection. Mice were fed a MAC-deficient (MD) diet or diets containing inulin or FOS as the sole MAC source and were subjected to murine model CDI. Burdens of *C. difficile* in mouse feces were quantified until 19 days post infection and are shown as blue circles for mice fed the MD diet, purple triangles for mice fed the FOS-containing diet, and red squares for the inulin-containing diet. The geometric mean of *C. difficile* burdens for mice fed each diet are connected with lines matching this diet-specific coloring scheme. The limit of detection of the *C. difficile* quantification assay (20,000 cfu *C. difficile*/mL feces) is shown as a horizontal dotted black line.

To begin to understand the differential impacts of these two MAC types on CDI, we grew *C. difficile* 630 in minimal medium supplemented with FOS or inulin. *C. difficile* 630 grows to a higher density in minimal medium supplemented with FOS relative to minimal medium supplemented with inulin (**Figure 2A**). This and previous work demonstrate that *C. difficile* cannot use inulin for growth [31]. However, *C. difficile* encodes an uncharacterized carbohydrate-active enzyme (CAZYme) that belongs to glycoside hydrolase family 32 (GH32), encoded by CD630_18050 in *C. difficile* 630. GH32 CAZYmes are important for fructan hydrolysis and are highly specific for their substrates (e.g. inulin, FOS, levan, sucrose) [32]. Together, these observations led us to hypothesize that FOS does not suppress CDI because *C. difficile* metabolizes FOS via a FOS-specific GH32 enzyme allowing it to persist during infection in mice fed FOS. To address this hypothesis, we performed high performance anion exchange chromatography with pulsed amperometric detection (HPAEC-PAD) to determine the extent of FOS utilization by *C. difficile* grown in FOS-supplemented minimal medium. We determined that *C. difficile* does not utilize FOS but instead consumes the trace amounts of glucose and fructose in the FOS preparation (**Figure 2B**, peaks within gray bars correspond to glucose and fructose based on reference chromatograms in **Figure S1**). Therefore, this work supports previous findings that *C. difficile* does not readily consume MACs [31] and that it is likely that factors unrelated to FOS metabolism by *C. difficile* contribute to the inability of FOS to clear murine CDI.

**Figure 2.**
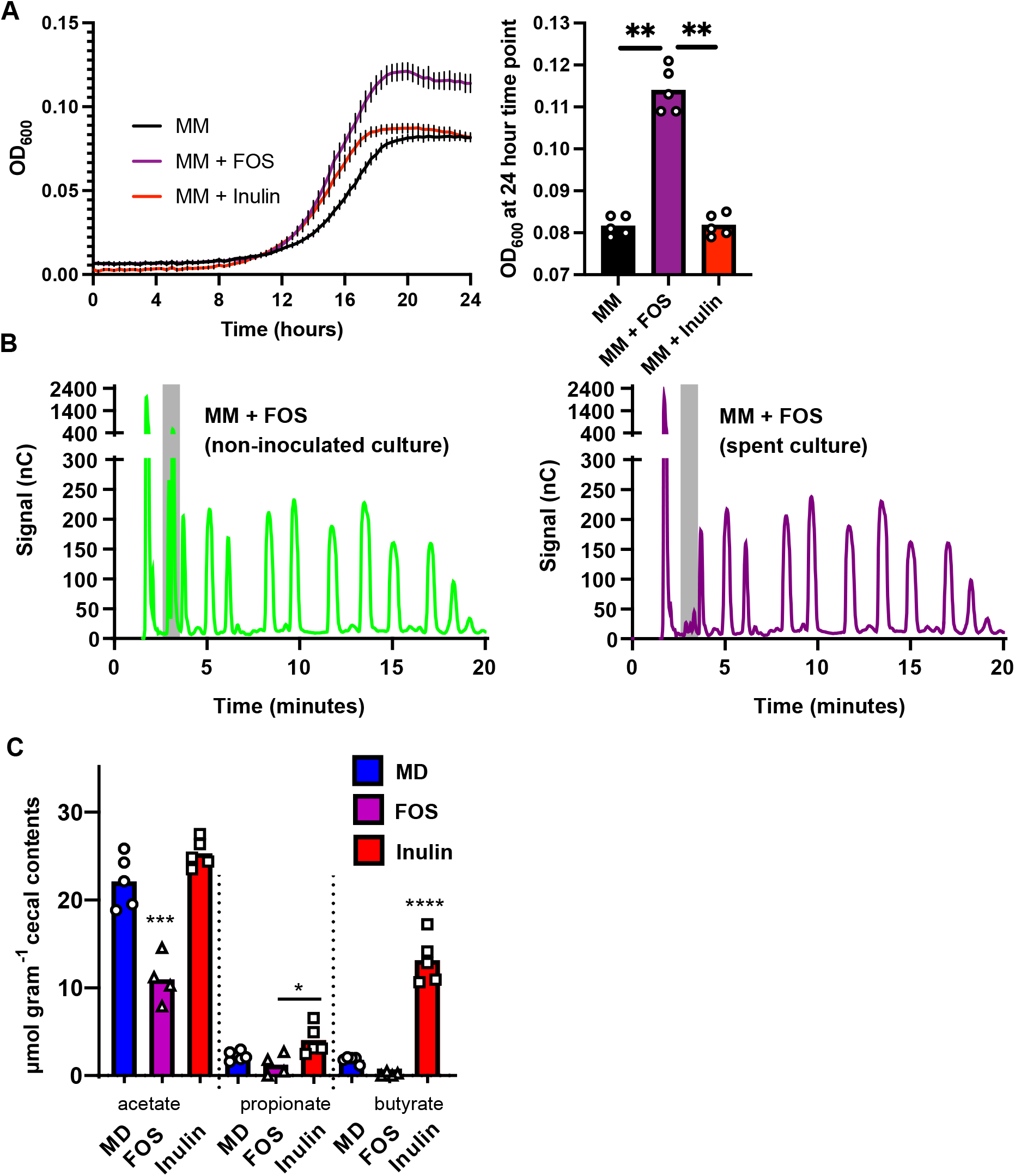
Growth of *C. difficile* 630 on FOS in vitro, and differential impacts of FOS and inulin on SCFA production by the microbiome in *C. difficile* 630-infected mice. (A) *C. difficile* 630 was grown in PETC-F minimal medium (MM, black lines), MM+FOS (purple lines), and MM+Inulin, red lines (n=5 biological replicates per strain) for 24 hours and culture density (OD_600_) was monitored (left). Lines and error bars represent mean OD_600_ readings and standard deviation at each time point, respectively. Statistically significant differences between the final OD_600_ of these cultures was determined by Mann-Whitney test (right). (B) Filtered supernatants from these cultures were analyzed using high performance anion exchange chromatography and a pulsed amphoteric detector (HPAEC-PAD). Representative chromatograms are shown for uninoculated media (left, green) and for spent media (right, purple). See **Figure S1** for a reference chromatogram, demonstrating that the metabolites depleted by *C. difficile* are monosaccharides that contaminate the FOS preparation rather than FOS itself. (C) The SCFAs acetate, propionate, and butyrate were quantified in the cecal contents of mice described in **Figure 1**, collected after euthanasia at 19 days post-infection. Individual data points represent SCFA concentrations measured via GC-MS and bars represent mean concentration. Blue bars represent SCFAs quantified in mice fed the MAC deficient diet, purple bars represent SCFAs quantified in mice fed the FOS-containing diet, and red bars represent SCFAs quantified in mice fed the inulin-containing diet. Statistical significance was determined by one-way ANOVA with Tukey’s multiple comparison test. See also **Figure S1**

The major metabolic end products of MAC metabolism by the gut microbiome are SCFAs, predominantly acetate, propionate, and butyrate [17,33,34]. Based on the metabolic capabilities of a given microbiome, MACs can differentially impact SCFA abundance and ratios in the gut. Our previous work and the work of others showed that SCFAs influence the fitness of *C. difficile* in animal models and in culture [8,19,28,35] and that FOS and inulin differentially impact the quantities and proportions of SCFAs produced by gut microbes in vitro [36]. We therefore hypothesized that FOS and inulin differentially impact CDI based on the quantities and types of SCFAs produced by the microbiome during infection. To address this hypothesis, we quantified acetate, propionate, and butyrate in the cecal contents of conventional mice fed FOS, inulin, or a MAC-deficient diet (see **Figure 1**) as described previously [8]. Mice fed FOS have lower levels of acetate, propionate, and butyrate in their ceca relative to those fed inulin (**Figure 2C**). In addition, less acetate was detected in the cecal contents of FOS-fed mice relative to mice fed a MAC deficient diet (**Figure 2C**), suggesting that alternative metabolic end products, distinct from acetate, propionate, and butyrate, are produced by FOS-fed microbiomes in this model. Consistent with our previous work, these data suggest that MACs that favor a SCFA-enriched gut environment discourage CDI.

### Cecal butyrate concentrations differentiate mice that do and do not suppress CDI across diverse MAC types

The conclusions that elevated SCFAs negatively impact *C. difficile* burdens in the mouse gut are based on experiments that used a limited number of dietary conditions. Specifically, both a complex MAC-rich diet (5010 Purina LabDiet) and a diet containing inulin as the sole MAC source suppress CDI. On the other hand, MAC deficient diets or a diet containing FOS as the sole MAC source do not clear CDI (see **Figure 1** and [8]). To further generalize these findings, we fed 5 additional diets containing different MAC sources to mice with experimental CDI. These diets contained one of three individual human milk oligosaccharides (HMOs; 2′-fucosyllactose (2′-FL), 6′-siaylyllactose (6′-SL), lacto-N-neo-Tetraose (LNnT)), a digestion resistant maltodextrin, or a complex mixture of MACs found within gum arabic. These MACs were selected based on evidence that HMOs impact SCFA production by gut microbes [37] and have a variety of beneficial effects on the eukaryotic host [38] and to understand whether the SCFAs produced by other structurally unrelated plant polysaccharides (distinct from fructans or the complex mixture of MACs present in standard rodent diets) impact *C. difficile* infection. We observed that these MAC types differentially impact *C. difficile* burdens and that out of these additional MACs tested, maltodextrin was the only one that consistently reduced *C. difficile* burdens below detection (**Figure 3A**). We then quantified acetate, propionate, and butyrate in the cecal contents of mice shown in **Figure 3A** to determine if SCFA concentrations differentiate mice with and without detectable fecal *C. difficile* in this cohort of mice fed inulin, gum arabic, resistant maltodextrin, 6′-SL, 2′-FL, and LNnT (**Figure 3B**). This diet-agnostic analysis of SCFA levels demonstrates that mice that cleared *C. difficile* below detection have significantly elevated levels of butyrate (but not acetate or propionate) in their cecal contents relative to mice with detectable *C. difficile*.

**Figure 3.**
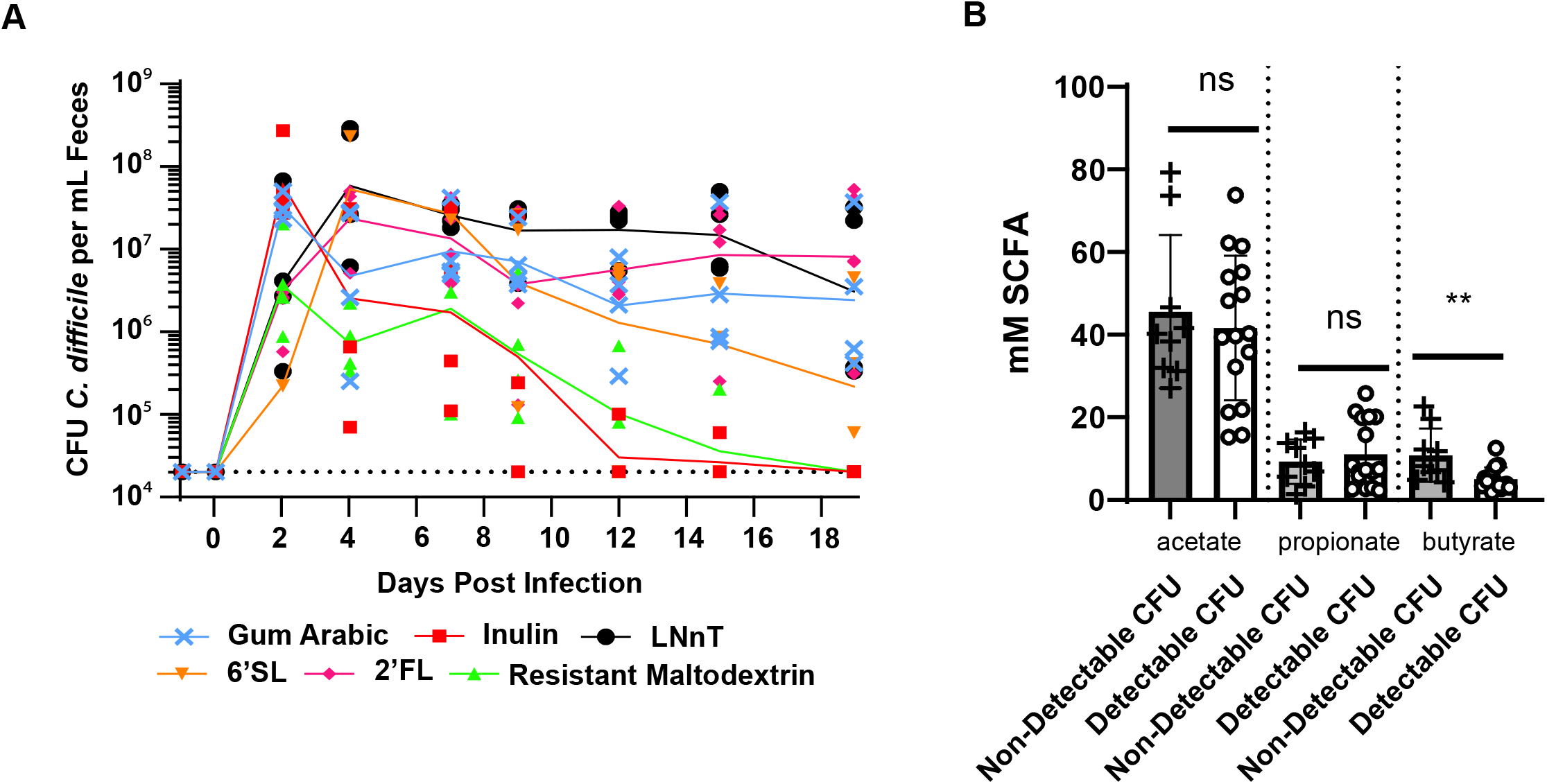
Differential impacts of MACs on *C. difficile* burdens and an association of *C. difficile* 630 clearance with cecal butyrate concentrations in mice. (A) Burdens of *C. difficile* in mouse feces were quantified until 19 days post infection and are shown as light blue crosses for mice fed the gum arabic diet, red squares for the inulin-containing diet, light green upward triangles for the resistant maltodextrin diet, orange downward triangles for the 6′-SL diet, magenta diamonds for the 2′-FL diet, and black circles for the LNnT diet. The geometric means of *C. difficile* burdens for mice fed each diet are connected with lines matching this diet-specific coloring scheme. The limit of detection is displayed as a dotted line. (B) The SCFAs acetate, propionate, and butyrate were quantified in cecal contents collected from mice after euthanasia at 19 days post infection via LC/MS. Individual measurements are shown as circles, squares, and triangles (acetate, propionate, and butyrate, respectively) and are stratified by mice that had detectable *C. difficile* in their feces versus those that had undetectable *C. difficile* in their feces. Means are displayed as bars and statistical significance was assessed by Mann-Whitney test.

### Fecal butyrate concentrations differentiate stool samples from humans with and without CDI

After learning that butyrate concentrations differentiate mice with and without detectable *C. difficile* in their feces, we wanted to know if butyrate concentrations are similarly associated with CDI in humans. Though previous work showed that SCFA concentrations increase in stool from CDI patients after a fecal transplant [39], the differences in concentrations of SCFAs in humans with CDI versus healthy controls was not previously determined. We quantified acetate, propionate, and butyrate in stool samples collected from patients who received care at Stanford Hospital in 2015. These stool samples were from patients with symptomatic CDI (diarrhea and positive for CDI (via Cepheid Xpert *C. difficile*)) and patients without CDI (negative for CDI (via Cepheid Xpert *C. difficile*)). In stool from the symptomatic *C. difficile* patients, we observed significantly lower concentrations of butyrate (but not acetate or propionate) relative to patients without CDI (**Figure 4**), which demonstrates that our findings in mice (**Figure 3B**) are generalizable to humans with CDI. Though acetate, propionate, and butyrate were previously shown to negatively impact the fitness of C. difficile and other bacterial pathogens [8,40], our observations from mice and humans provide the rationale for focused and specific investigation of butyrate.

**Figure 4.**
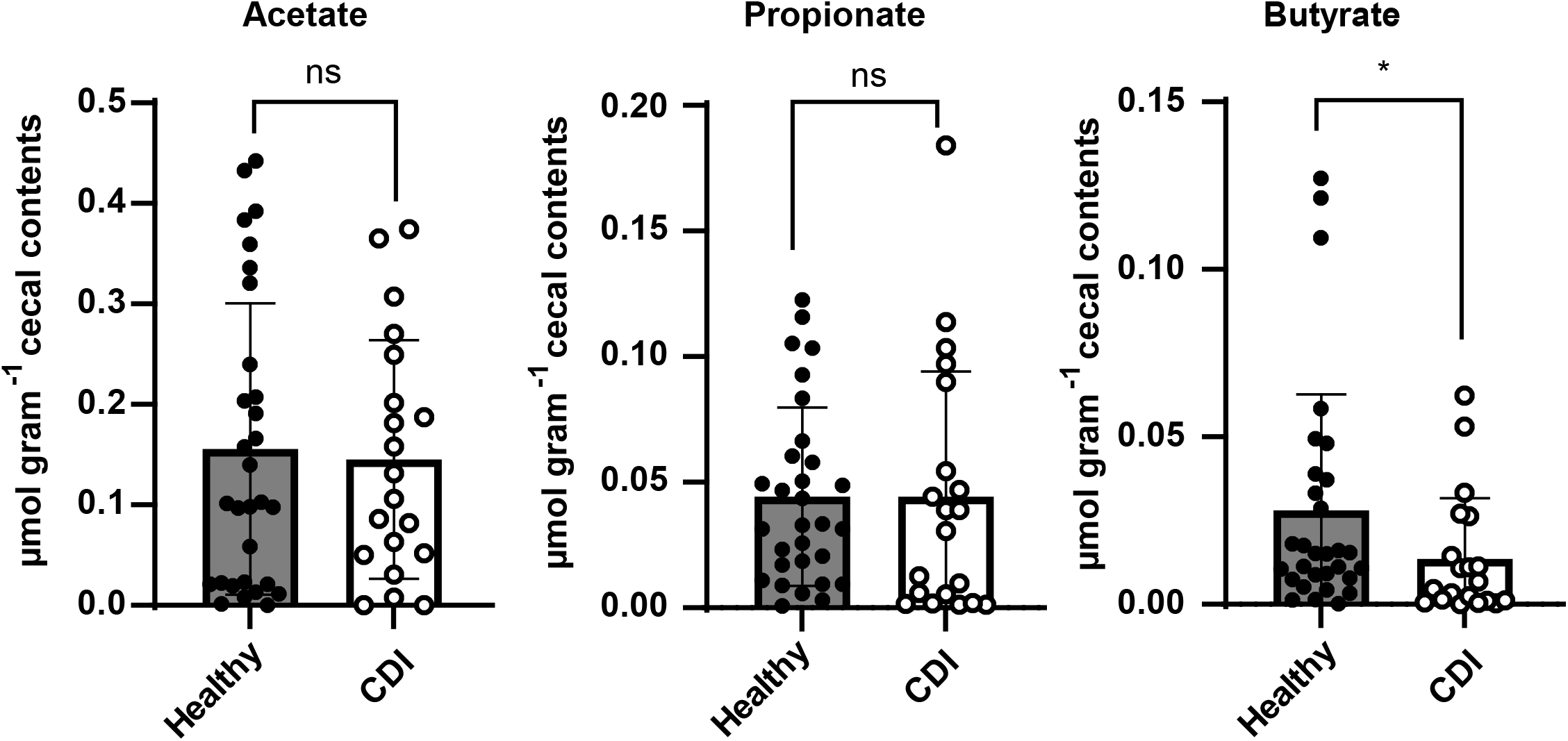
Fecal butyrate concentrations differentiate humans with CDI from healthy controls. The SCFAs acetate, propionate, and butyrate were quantified in human stool samples from patients with symptomatic CDI and from healthy controls via GC-MS (see **Human subjects/patient enrollment)**. Means are displayed as bars and statistically significant differences in SCFA concentrations between each patient population was determined by Mann-Whitney test.

### Butyrate negatively impacts growth in diverse C. difficile isolates

Our previous work showing that butyrate impacts *C. difficile* growth was restricted to the commonly studied *C. difficile* 630 strain [8] and **Figures 1-3**). Though similar butyrate-dependent effects were observed in 4 unsequenced *C. difficile* isolates [18], we sought to further situate these findings in the context of a phylogenetically diverse sample of *C. difficile* strains. We grew 13 different *C. difficile* isolates with representatives from 10 ribotypes (including *C. difficile* 630; **Table 1**) in pure culture in the presence of 0, 6.25, 12.5, 25, and 50mM sodium butyrate and in matched concentrations of sodium chloride. For all *C. difficile* strains tested, butyrate negatively impacts growth kinetics (**Figure S2**), with notable concentration-dependent differences in maximum growth rate (**Figure 5A**) and lag time (**Figure 5B**). All strains tested had significantly longer lag times in the presence of 50mM butyrate compared to 0mM butyrate. Similarly, all but 2 strains tested (CD196 and TL178) exhibited significantly reduced maximum growth rates in the presence of 50mM butyrate compared to 0mM butyrate. The significance and magnitude of these effects were smaller for intermediate butyrate concentrations but were concentration-dependent.

**Table 1:**
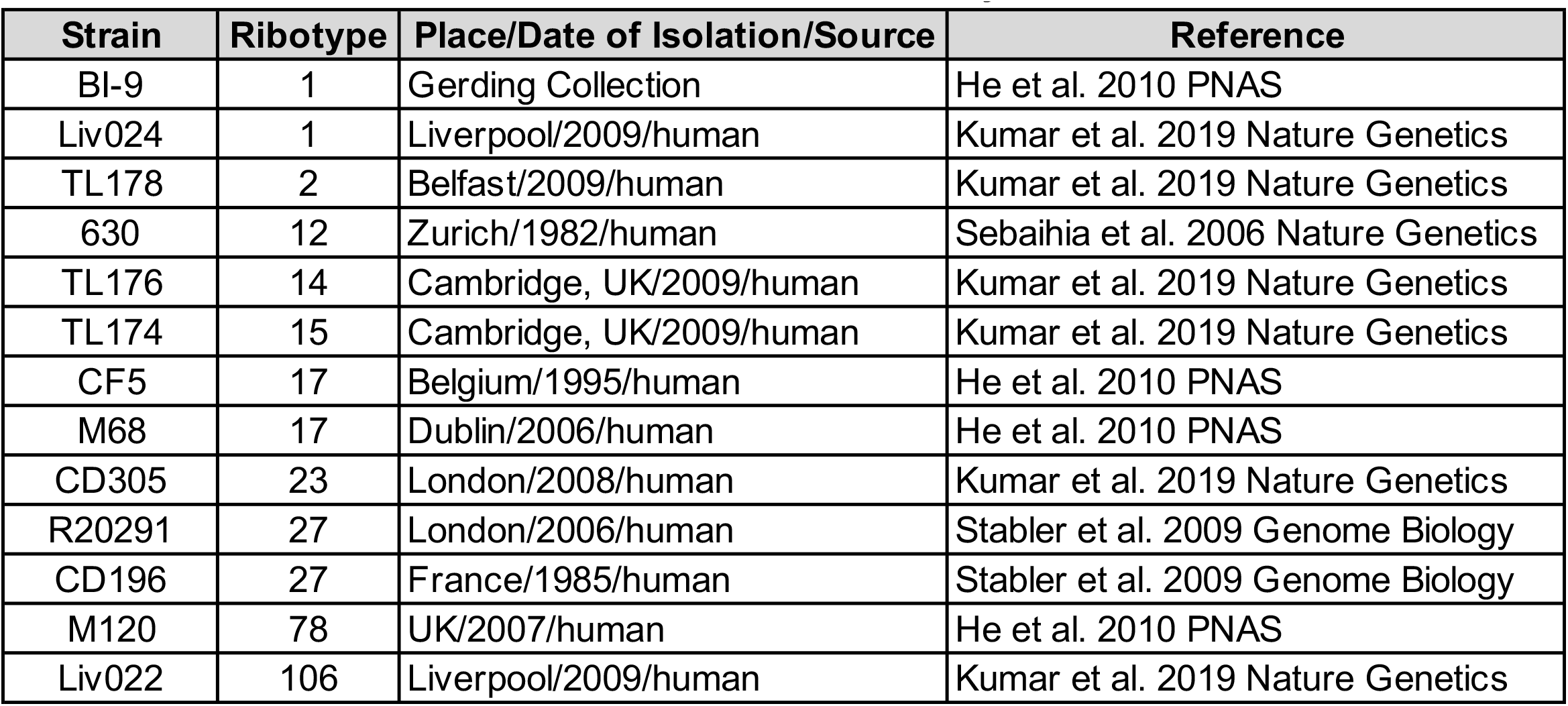
*Clostridioides difficile* strains used in this study.

| **Strain** | **Ribotype** | **Place/Date of Isolation/Source** | **Reference** |
| --- | --- | --- | --- |
| BI-9 | 1 | Gerding Collection | He et al. 2010 PNAS |
| Liv024 | 1 | Liverpool/2009/human | Kumar et al. 2019 Nature Genetics |
| TL178 | 2 | Belfast/2009/human | Kumar et al. 2019 Nature Genetics |
| 630 | 12 | Zurich/1982/human | Sebahia et al. 2006 Nature Genetics |
| TL176 | 14 | Cambridge, UK/2009/human | Kumar et al. 2019 Nature Genetics |
| TL174 | 15 | Cambridge, UK/2009/human | Kumar et al. 2019 Nature Genetics |
| CF5 | 17 | Belgium/1995/human | He et al. 2010 PNAS |
| M68 | 17 | Dublin/2006/human | He et al. 2010 PNAS |
| CD305 | 23 | London/2008/human | Kumar et al. 2019 Nature Genetics |
| R20291 | 27 | London/2006/human | Stabler et al. 2009 Genome Biology |
| CD196 | 27 | France/1985/human | Stabler et al. 2009 Genome Biology |
| M120 | 78 | UK/2007/human | He et al. 2010 PNAS |
| Liv022 | 106 | Liverpool/2009/human | Kumar et al. 2019 Nature Genetics |

**Figure 5.**
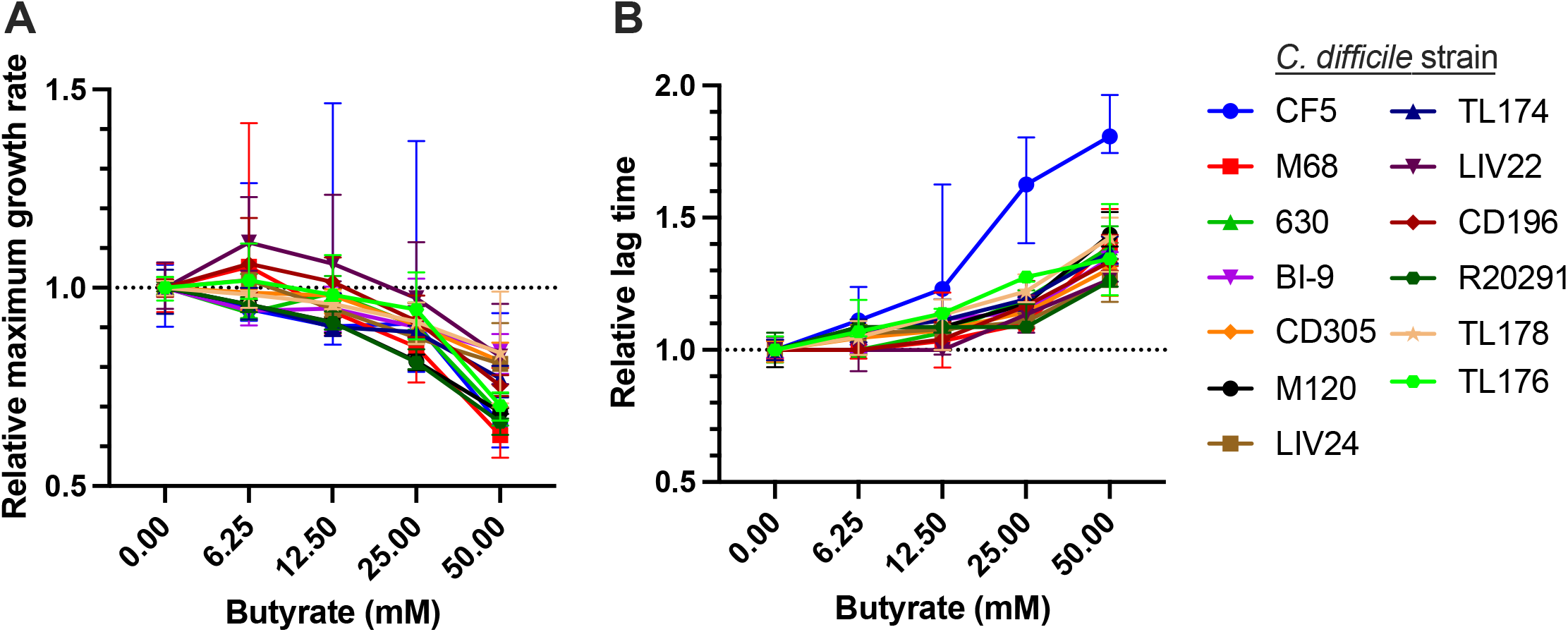
Butyrate negatively impacts growth in diverse *C. difficile* strains. Thirteen *C. difficile* strains (see **Table 1**) were grown anaerobically in mRCM supplemented either with 0, 6.25, 12.5, 25, or 50mM sodium butyrate or matched concentrations of sodium chloride (NaCl) for 24 hours. Culture density (OD_600_) was monitored throughout this time course (n=6 replicates per growth condition per strain). For all cultures (A) maximum growth rate and (B) lag time were calculated. All strains tested had significantly longer lag times in the presence of 50mM butyrate compared to 0mM butyrate. Similarly, all but 2 strains tested (CD196 and TL178) exhibited significantly reduced maximum growth rates in the presence of 50mM butyrate compared to 0mM butyrate. Median line is displayed and statistically significant differences between relevant groups were determined by Mann-Whitney test. **Figure S2** shows representative growth curves for all 13 strains under the growth conditions tested.

Though bulk measurements of butyrate in human and mouse samples are lower than 50 mM (**Figure 2, Figure 3, Figure 4**, and [8,39], concentrations of butyrate produced by microbiome members in the gut at relevant spatial scales (e.g., when *C. difficile* is in close proximity to butyrate-producing commensals) remains unclear but is likely higher than what is observed via bulk measurements. Regardless, the concentration-dependent effects we observe for all strains (**Figure 5**) demonstrate that *C. difficile* growth is reliably impacted by butyrate and suggest that the molecular mechanisms underlying this response are conserved across diverse *C. difficile* strains.

## DISCUSSION

This work adds to the growing body of literature that demonstrates that diet impacts CDI in animal models of infection. Specifically, it refines previous observations about the impacts of MACs on *C. difficile* fitness in the gut by showing that diets which lead to elevated butyrate production by the microbiome reduce burdens of *C. difficile* during infection. Taken together, our work and the work of others shows that inulin, maltodextrin, and xanthan gum are purified MACs that consistently suppress CDI while FOS, 2′-FL, 6′-SL, and LNnT are purified MACs that do not suppress CDI (**Figure 3**, [8,9]). Unlike a standard rodent diet that is a complex mixture of MACs [8], we show that a different complex mixture of MACs (gum arabic) does not suppress *C. difficile* burdens in mice (**Figure 3**). Importantly, given that our work exclusively used conventionally-reared Swiss-Webster mice and that differences in microbiome configuration dictate metabolites used and produced by a given community [41], it is possible that the MAC sources that did not clear CDI in our model would clear CDI in the context of a different microbiome or host. As such, future work should consider the variability of microbiome composition and metabolic outputs when designing dietary strategies for impacting CDI and other disease states.

*C. difficile* burdens are unlikely to be the only parameter impacted by MACs during infection, which highlights additional directions for future work. For example, though we observed that FOS does not suppress CDI in mice (**Figure 1**), it was previously shown that FOS increases survival time in hamsters infected with *C. difficile* [10] but the mechanism of this protection was not defined. As these and other MAC-driven impacts on the host immune system are better understood, they will likely contribute to the formulation of specific diet-based strategies to simultaneously bolster the host immune response while reducing the fitness of *C. difficile*, either through the manipulation of SCFA levels (which influence inflammation [42] and colonocyte metabolism [43,44] or by directly impacting the mucosal immune system (e.g., via HMOs which can influence inflammatory cell populations [45] and positively impact barrier function [46]. Future diet-based strategies to mitigate CDI will similarly be informed by the growing literature surrounding the impact of other dietary inputs on CDI (see **Introduction**).

Because butyrate levels differentiate mice and humans that have CDI from those that do not (**Figure 2C, 3B, 4**), continued focus on this SCFA in the context of CDI will yield important insights into the biology of *C. difficile*, the ecology of CDI, and future therapeutic approaches. We and others previously showed that butyrate negatively impacts growth in 5 distinct *C. difficile* strains [8,18] and in the current study we extend these findings to 12 additional *C. difficile* strains (**Figure 5, Table 1**), together demonstrating that these phenotypes are generalizable across a large sample of *C. difficile* clinical isolates. We recently developed a conceptual model to unify the seemingly paradoxical observations that growth and toxin production are differentially impacted by butyrate [35]. Specifically, *C. difficile* infection and proliferation is favored in a dysbiotic (butyrate deficient) gut environment where there is minimal competition for metabolites (e.g., amino acids, organic acids, sugars). Under these conditions, *C. difficile* produces no detectable toxin. However, as the microbiome recovers from dysbiosis, the availability of metabolites decreases and the concentrations of butyrate increases, resulting in reduced *C. difficile* fitness. In response to these conditions, *C. difficile* up-regulates its toxins, which increase inflammation [47], and presumably helps to re-establish facets of microbiome community function that allow *C. difficile* to thrive.

Future work based on the above conceptual model and the data presented in the current study will seek to understand the variety of host-by-microbiome-by-diet interactions that influence *C. difficile* fitness in the gut. Specific foci on the molecular mechanisms and genetic circuitry underlying the responses of *C. difficile* to butyrate will facilitate a better basic understanding of *C. difficile* and how it interacts with the host and the gut microbiome. In addition, continued research on these and other diet-driven effects on CDI are likely to yield insights that will aid in the development of specific and targeted manipulation of CDI, either through dietary intervention, therapeutic application of specific microbes (e.g., probiotics), or delivery of specific metabolites.

## METHODS

### Bacterial strains and culture conditions

Frozen stocks of *C. difficile* strains used in the study (**Table 1;** [48–50]) were maintained as -80°C stocks in 25% glycerol under anaerobic conditions in septum-topped vials. *C. difficile* was routinely cultured on CDMN agar, composed of *C. difficile* agar base (Oxoid) supplemented with 7% defibrinated horse blood (HemoStat Laboratories), 32 mg/L moxalactam (Santa Cruz Biotechnology), and 12 mg/L norfloxacin (Sigma-Aldrich) in an anaerobic chamber at 37° (Coy).

After 16-24 hours of growth, a single colony was picked into 5 mL of pre-reduced reinforced clostridial medium (RCM, Oxoid), modified reinforced Clostridial medium (mRCM: 10g/L beef extract, 3g/L yeast extract, 10g/L peptone, 5g/L dextrose, 5g/L sodium chloride, 3g/L sodium acetate, 0.5g/L cysteine hydrochloride) or PETC medium (ATCC medium 1754) without fructose (PETC-F), and grown anaerobically at 37°C for 16-24 hours. Liquid cultures were used as inocula for growth curves and for experiments using murine model CDI, below.

For in vitro growth curve experiments examining *C. difficile* fructan utilization, subcultures were prepared at a 1:200 dilution in pre-reduced PETC-F minimal medium supplemented with either 5 mg/mL inulin (OraftiHP; Beneo-Orafti group) or 5 mg/mL FOS (Orafti P95, Beneo-Orafti group) in sterile polystyrene 96 well tissue culture plates with low evaporation lids (Falcon).

Cultures were grown anaerobically as above in a BioTek Powerwave plate reader. At 15 minute intervals, the plate was shaken on the ‘slow’ setting for 1 minute and the optical density (OD_600_) of the cultures was recorded using Gen5 software (version 1.11.5). After 24 hours of growth, culture supernatants were collected, centrifuged (5 minutes at 2,500 x g), filtered (0.22 µm PVDF filter), and stored at -20°C for high performance anion exchange chromatography, below.

For in vitro growth curve experiments examining *C. difficile* growth in the presence of butyrate, subcultures were prepared at a 1:200 dilution in pre-reduced mRCM (RCM lacking starch and agar which reduces clumping artefacts in OD_600_ readings) in sterile polystyrene 96 well tissue culture plates with low evaporation lids (Falcon). Cultures were grown anaerobically in a BioTek Epoch2 plate reader. At 30-minute intervals the plate was shaken on the ‘slow’ setting for 1 minute and the OD_600_ of the cultures was recorded using Gen5 software (version 1.11.5).

### Murine model of C. difficile infection

All animal studies were conducted in strict accordance with Stanford University Institutional Animal Care and Use Committee (IACUC) guidelines. Murine model CDI was performed on age- and sex-matched conventionally-reared Swiss-Webster mice (Taconic) between 8 and 17 weeks of age.

To reduce colonization resistance against *C. difficile*, mice were given a single dose of clindamycin by oral gavage (1 mg/mouse; 200 µL of a 5 mg/mL solution) and were infected 24 hours later with 200 µL of overnight culture grown in RCM (approximately 1.5×10^7^ cfu/mL).

Feces were collected from mice directly into microcentrifuge tubes and immediately placed on ice. To monitor *C. difficile* burdens in feces, 1 µL of each fecal sample was resuspended in PBS to a final volume of 200 µL, 10-fold serial dilutions of fecal slurries (through 10^−3^-fold) were prepared in sterile polystyrene 96 well tissue culture plates (Falcon). For each sample, two 10 µL aliquots of each dilution (technical replicates) were spread onto CDMN agar supplemented with erythromycin (100 mg/L, Acros Organics). Erythromycin supplementation further reduces growth of bacteria from mouse feces and has no impact on *C. difficile* colony counts (data not shown). After 16–24 hours of anaerobic growth at 37°C, colonies were enumerated and technical replicates were averaged to determine *C. difficile* burdens in feces (limit of detection = 2×10^4^ cfu/mL feces). Immediately following euthanasia at 19 days post infection, cecal contents were removed from mice, weighed, and flash frozen in liquid nitrogen. *C. difficile* was undetectable in all mice prior to inoculation with CDI.

### Mouse diets

Mice were fed one of eight custom diets (Bio Serv) ad libitum: (1) a MAC-deficient control diet containing 68% glucose (w/v), 18% protein (w/v), and 7% fat (w/v) (MD, Bio-Serv); or diets containing 10% (w/v) of one of the following ingredients as a sole source of MAC: (2) inulin (Orafti HP; Beneo-Orafti group, Mannheim, Germany), (3) FOS (Orafti P95, Beneo-Orafti group, Mannheim, Germany), (4) gum arabic (Nutriloid Gum Arabic FT; TIC Gums, Belcamp, Maryland), (5) digestion resistant maltodextrin (Fibersol-2; ADM/Matsutani LLC, Chicago, Illinois), (6) lacto-N-neotetraose (LNnT; Kyowa Hakko, Tokyo, Japan), (7) 2′-fucosyllactose (2′-FL; Inalco SpA, Milano, Italy), or (8) 6′-sialyllactose (6′-SL; Inalco SpA, Milano, Italy). HMOs were enzymatically (LNnT) or chemically synthesized (2′-FL, 6′-SL). For MAC-containing diets, MAC ingredients were swapped for an equal quantity of glucose.

### Human subjects/patient enrollment

Human stool samples were collected from patients receiving care at Stanford Health Care between January 2015 and November 2015 and participating in an IRB-exempt quality improvement project aimed at understanding the rates of *C. difficile* transmission in hematopoietic stem cell transplant patients. Samples are either from the patient’s first post-admission bowel movement or were collected at a frequency no more than once every 7 days post admission. Samples were collected and immediately assayed for *C. difficile* TcdB using the Xpert *C. difficile* assay (Cepheid). Patients with unformed, *C. difficile*+ stools, were considered to have CDI. After this diagnostic procedure, residual de-identified samples (regardless of CDI status) were stored at 4°C for no more than 48 hours and frozen at -80°C. Samples were subjected to targeted metabolomics, where the SCFAs acetate, propionate, and butyrate were quantified (see **SCFA quantification**, below).

### Quantification of FOS-degradation products

To quantify FOS degradation by *C. difficile*, spent and non-inoculated PETC-F medium supplemented with 5 mg/mL FOS were filtered through 0.22 µm PVDF filters, dialysed through centrifuge filters (10 kDa MWCO, Millipore) and diluted with deionized water to bring the concentration of carbohydrate sources to a concentration of 1 µg/µL except for inulin (10 µg/µL). Samples were subjected to high performance anion exchange chromatography on a Dionex ICS-5000 system with an AS-AP autosampler and a pulsed amperometric detector, using a Dionex CarboPak PA1 column (4×250 mm Analytical, Thermo Scientific) with a corresponding 4×50 mm guard column. The following solvent gradient was used (A = 100 mM NaOH, B = 100 mM NaOH 1 M NaOAc): 0 to 60 minutes, 5% to 45% B; 60 to 70 minutes, 45% to 75% B. To prepare the reference chromatograms shown in **Figure S1**, individual 5 mg/mL solutions of fructose, glucose, sucrose, kestose, nystose, and FOS were prepared in distilled water, filtered through 0.22 µm PVDF filters, and subjected to HPAEC-PAD as described above.

### SCFA quantification

Two methods were used to quantify SCFAs in cecal contents from mice and in human stool: (1) a GC-MS-based method used in our previous work [8] and (2) an LC-MS-based method developed to overcome restrictions to access of core facility equipment during the early stages of the COVID-19 pandemic at Stanford University.

*GC-MS-based SCFA quantification*. Cecal contents from mice or human stool (70-150 mg) were suspended in a final volume of 600 µl in ice-cold ultra-pure water and blended with a pellet pestle (Kimble Chase) on ice. The slurry was centrifuged at 2,350 × g for 30 seconds at 4°C and 250 µL of the supernatant was removed to a septum-topped glass vial and acidified with 20µL HPLC grade 37% HCl (Sigma Aldrich). Diethyl ether (500 µL) was added to the acidified cecal supernatant to extract SCFAs. Samples were then vortexed at 4°C for 20 minutes on ‘high’ and then were centrifuged at 1,000 × g for 3 minutes. The organic phase was removed into a fresh septum-topped vial and placed on ice. Then, a second extraction was performed with diethyl ether as above. The first and second extractions were combined for each sample and 250 µL of this combined solution was added to a 300 µL glass insert in a fresh glass septum-topped vial containing and the SCFAs were derivatized using 25 µL N-tert-butyldimethylsilyl-N-methyltrifluoroacetamide (MTBSTFA; Sigma Aldrich) at 60°C for 30 minutes.

Analyses were carried out using an Agilent 7890/5975 single quadrupole GC/MS. Using a 7683B autosampler, 1 µL split injections (1:100) were made onto a DB-5MSUI capillary column (30 m length, 0.25 mm ID, 0.25 µm film thickness; Agilent) using helium as the carrier gas (1 mL/minute, constant flow mode). Inlet temperature was 200°C and transfer line temperature was 300°C. GC temperature was held at 60°C for 2 minutes, ramped at 40°C/min to 160°C, then ramped at 80°/min to 320°C and held for 2 minutes; total run time was 8.5 minutes. The mass spectrometer used electron ionization (70eV) and scan range was m/z 50–400, with a 3.75-minute solvent delay. Acetate, propionate, and butyrate standards (20 mM, 2 mM, 0.2 mM, 0.02 mM, 0 mM) were acidified, extracted, and derivatized as above, were included in each run, and were used to generate standard curves to enable SCFA quantification.

*LC-MS-based SCFA quantification*. The LC-MS-based SCFA quantification method was adapted from [51]. Briefly, cecal contents from mice (50 to 150 mg) were weighed on an analytical balance and diluted in extraction buffer containing: 80% HPLC-grade water (Fisher), 20% HPLC-grade acetonitrile (ACN; Fisher) and labelled isotopes of each SCFA measured (2.5 uM d3-acetic acid (Sigma Aldrich), 1 uM propionic-3,3,3-d3 acid (CDN Isotopes), 0.5 uM butyric-4,4,4-d3 acid (CDN Isotopes)). The volume of extraction buffer in microliters was 4X the mass of cecal contents in milligrams for each sample.

Acid-washed beads (150uM-212uM; SigmaAldrich G1145-10G) were added to the samples and the samples were shaken at 30 Hz for 10 minutes to homogenize and extract the metabolites. The samples were then incubated at -20°C for 1 hour and subsequently centrifuged at 4°C for 5 minutes at 12,000 rcf. 40 uL of the supernatant was transferred to a 96 well plate to which 20 uL of 200mM 3-nitrophenylhydrazine hydrochloride (Sigma Aldrich; dissolved in 50% ACN and 50% water) and 20 uL of 120 mM 1-ethyl-3-(3-dimethylaminopropyl)carbodiimide hydrochloride (Pierce; dissolved in 47% ACN, 47% water and 6% HPLC-grade pyridine (Sigma Aldrich)) were added. The plate was then sealed and shaken in an incubator at 37°C for 30 minutes. After 30 minutes the plate was cooled to 4°C and 20 uL of the reaction volume was transferred to 980 μL of a 90:10 (v/v) Water:ACN solution.

Analyses were carried out using an Agilent 6470 triple quadrupole LC/MS. Using a G7167B multisampler, 10uL injections were made onto an Acquity UPLC BEH C18 column (100 mm length, 2.1 mm inner diameter, 130 Å pore size, 1.7 um particle size; Waters) using water:formic acid (100:0.01, v/v; solvent A) and acetonitrile:formic acid (100:0.01, v/v; solvent B) as the mobile phase for gradient elution. The column flow rate was 0.35 mL/min; the column temperature was 40°C, and the autosampler was kept at 5°C. The binary solvent elution gradient was optimized at 15% B for 2 min, 15%–55% B in 9 min, and then held at 100% B for 1 min. The column was equilibrated for 3 min at 15% B between injections. The drying gas (N2) temperature was set to 300°C with a flow rate of 12 L/min. The sheath gas temperature was also set to 300°C with a flow rate of 12 L/min. The nebulizer gas was set to 25 PSI and the capillary voltage was set to 4200 V.

Quantification of analytes was done by standard isotope dilution protocols. In brief, serial dilutions of a 3 SCFA standard solution (10 mM, 1 mM, 0.1 mM, 0.01 mM, 0.001 mM, and 0 mM) were derivatized as above and included in each run to verify sample concentrations were within linear ranges. For samples within linear range, analyte concentration was calculated as the product of the paired internal standard concentration and the ratio of analyte peak area to internal standard peak area. A single product ion was used for each analyte, no secondary or qualifier ions were used. To ensure the highest signal-to-noise ratio, the following steps were taken. First, to ensure that the predicted singly derivatized species was the dominant precursor ion, full-mass Q1 scans were performed over the m/z range 100 to 300. Second, collision energies and fragmentor voltage were optimized using Agilent’s MassHunter Optimizer program with direct infusion of the derivatives from individual standard solutions containing 50mM of each fatty acid. Optimizer was set to search collision energies from -10V to -120V in 10V increments and select the two most intense product ions for optimization. Fragmentor voltage had minimal impact and was manually set to 75 V.

### Measurement of maximum growth rate and lag time for in vitro growth experiments

Raw OD_600_ measurements of cultures grown in mRCM (see ‘Bacterial strains and culture conditions’, above) were exported from Gen5 and analyzed using the growth_curve_statistics.py script (see **Code Availability**, below). Growth rates were determined for each culture by calculating the derivative of natural log-transformed OD_600_ measurements over time. Growth rate values at each time point were then smoothed using a moving average over 150-min intervals to minimize artefacts due to noise in OD measurement data, and these smooth growth rate values were used to determine the maximum growth rate for each culture. To mitigate any remaining issues with noise in growth rate values, all growth rate curves were also inspected manually. Specifically, in cases where the growth_curve_statistics.py script selected an artefactual maximum growth rate, the largest local maximum that did not correspond to noise was manually assigned as the maximum growth rate. Additionally, lag time was calculated as half the time to reach the maximum growth rate.

## Code availability

Python script that was used to compute maximum growth rate and lag time from growth curve data is freely available at https://github.com/HryckowianLab/Pensinger_2022.

## Statistical analysis

Statistical analysis was performed using Graphpad Prism 9.1.0. Details of specific analyses, including statistical tests used, are found in applicable figure legends. * = p<0.05, ** = p<0.005, *** = p<0.0005, **** = p<0.0001.

## Acknowledgments

We thank Keith Garleb (Abbott) for helpful comments and Niaz Banaei (Stanford University) for logistical assistance with human stool samples. This work was funded by Abbott (ZB40) and by grants from the following sources: NIH NIDDK (R01-DK085025 to JLS), NIH DCI (R01-CA200423 to CRB) an NIH postdoctoral NRSA (5T32AI007328 to AJH), a Stanford University School of Medicine Dean’s Postdoctoral Fellowship (AJH), a Feodor Lynen Postdoctoral Fellowship by the Alexander von Humboldt Foundation (BS), NSF Graduate Research Fellowships (WVT, CM), Howard Hughes Medical Institute (CRB), and startup funding from the University of Wisconsin-Madison (AJH). JLS received an Investigators in the Pathogenesis of Infectious Disease Award from the Burroughs Wellcome Fund and is a Chan Zuckerburg Biohub Investigator.

## Author Contributions

AJH, ATF, HAD, WVT, MMC, JOG, SKH, and BS performed experiments. AJH, ATF, HAD, MMC, WVT, JOG, CM, and DAP analyzed the data. DAP, AJH, HAD, JOG, and BS prepared the display items. EVR, CRB, JC, VA, RHB, LST, and JLS provided key insights, tools, and reagents. DAP and AJH wrote the paper.

All authors edited the manuscript prior to submission.

## Declaration of Interests

This work was funded in part by Abbott. This funder contributed to the design of the experiments shown in **Figure 3**.

## Figure Legends

**Table 1. Bacterial strains used in this study**. Related to **Figures 1, 2, 3, and 5.**

**Figure S1. HPAEC-PAD Chromatograms**. Reference chromatograms for fructose (light blue), glucose (orange), sucrose (gray), kestose (yellow), nystose (blue), and FOS (green) are shown. Chromatograms for FOS-supplemented PETC-F minimal medium (FOS + MM; dark blue) and spent PETC-F minimal medium (FOS + MM spent; peach) are also shown and duplicated from **Figure 2B**. Related to **Figure 2**.

**Figure S2. Representative growth curves of thirteen *C. difficile* strains grown in the presence of sodium butyrate and sodium chloride**. The thirteen *C. difficile* strains listed in **Table 1** were grown anaerobically in mRCM supplemented with either 0, 6.25, 12.5, 25, or 50mM sodium butyrate or identical concentrations of NaCl for 24 hours. Each plot shows three representative growth curves per strain per condition and represents raw culture density (OD_600_) measurements for each strain tested. Symbols represent mean and standard deviation of replicates. Related to **Figure 5**.

## REFERENCES

1. Centers for Disease Control and. Antibiotic resistance threats in the United States. US Dep Heal Hum Serv. 2019; 1–113. doi:10.15620/cdc:82532

2. Khanna S, Pardi DS, Aronson SL, Kammer PP, Orenstein R, St Sauver JL, et al. The epidemiology of community-acquired Clostridium difficile infection: a population-based study. Am J Gastroenterol. Am J Gastroenterol; 2012;107: 89–95. doi:10.1038/AJG.2011.398 PMID:22108454

3. David LA, Maurice CF, Carmody RN, Gootenberg DB, Button JE, Wolfe BE, et al. Diet rapidly and reproducibly alters the human gut microbiome. Nature. Nature; 2014;505: 559–563. doi:10.1038/NATURE12820 PMID:24336217

4. Wastyk HC, Fragiadakis GK, Perelman D, Dahan D, Merrill BD, Yu FB, et al. Gut-microbiota-targeted diets modulate human immune status. Cell. Cell; 2021;184: 4137-4153.e14. doi:10.1016/J.CELL.2021.06.019 PMID:34256014

5. Mefferd CC, Bhute SS, Phan JR, Villarama J V., Do DM, Alarcia S, et al. A High-Fat/High-Protein, Atkins-Type Diet Exacerbates Clostridioides (Clostridium) difficile Infection in Mice, whereas a High-Carbohydrate Diet Protects. mSystems. American Society for Microbiology; 2020;5. doi:10.1128/MSYSTEMS.00765-19/SUPPL_FILE/MSYSTEMS.00765-19-ST004.EPS

6. Moore JH, Pinheiro CCD, Zaenker EI, Bolick DT, Kolling GL, Van Opstal E, et al. Defined Nutrient Diets Alter Susceptibility to Clostridium difficile Associated Disease in a Murine Model. PLoS One. PLoS One; 2015;10. doi:10.1371/JOURNAL.PONE.0131829 PMID:26181795

7. Battaglioli EJ, Hale VL, Chen J, Jeraldo P, Ruiz-Mojica C, Schmidt BA, et al. Clostridioides difficile uses amino acids associated with gut microbial dysbiosis in a subset of patients with diarrhea. Sci Transl Med. Sci Transl Med; 2018;10. doi:10.1126/SCITRANSLMED.AAM7019 PMID:30355801

8. Hryckowian AJ, Van Treuren W, Smits SA, Davis NM, Gardner JO, Bouley DM, et al. Microbiota-accessible carbohydrates suppress Clostridium difficile infection in a murine model. Nat Microbiol. Nat Microbiol; 2018;3: 662–669. doi:10.1038/S41564-018-0150-6 PMID:29686297

9. Schnizlein MK, Vendrov KC, Edwards SJ, Martens EC, Young VB. Dietary Xanthan Gum Alters Antibiotic Efficacy against the Murine Gut Microbiota and Attenuates Clostridioides difficile Colonization. mSphere. mSphere; 2020;5. doi:10.1128/MSPHERE.00708-19 PMID:31915217

10. Wolf BW, Meulbroek JA, Jarvis KP, Wheeler KB, Garleb KA. Dietary Supplementation with Fructooligosaccharides Increase Survival Time in a Hamster Model of Clostridium difficile-Colitis. Biosci Microflora. JAPAN BIFIDUS FOUNDATION; 1997;16: 59–64. doi:10.12938/BIFIDUS1996.16.59

11. Collins J, Robinson C, Danhof H, Knetsch CW, Van Leeuwen HC, Lawley TD, et al. Dietary trehalose enhances virulence of epidemic Clostridium difficile. Nature. Nature; 2018;553: 291–294. doi:10.1038/NATURE25178 PMID:29310122

12. Eyre DW, Didelot X, Buckley AM, Freeman J, Moura IB, Crook DW, et al. Clostridium difficile trehalose metabolism variants are common and not associated with adverse patient outcomes when variably present in the same lineage. EBioMedicine. Elsevier; 2019;43: 347. doi:10.1016/J.EBIOM.2019.04.038 PMID:31036529

13. Zackular JP, Moore JL, Jordan AT, Juttukonda LJ, Noto MJ, Nicholson MR, et al. Dietary zinc alters the microbiota and decreases resistance to Clostridium difficile infection. Nat Med. Nat Med; 2016;22: 1330–1334. doi:10.1038/NM.4174 PMID:27668938

14. Rojo D, Gosalbes MJ, Ferrari R, Pérez-Cobas AE, Hernández E, Oltra R, et al. Clostridium difficile heterogeneously impacts intestinal community architecture but drives stable metabolome responses. ISME J. ISME J; 2015;9: 2206–2220. doi:10.1038/ISMEJ.2015.32 PMID:25756679

15. Jenior ML, Leslie JL, Young VB, Schloss PD. Clostridium difficile Colonizes Alternative Nutrient Niches during Infection across Distinct Murine Gut Microbiomes. mSystems. American Society for Microbiology; 2017;2. doi:10.1128/msystems.00063-17

16. Jenior ML, Leslie JL, Young VB, Schloss PD. Clostridium difficile Alters the Structure and Metabolism of Distinct Cecal Microbiomes during Initial Infection To Promote Sustained Colonization. mSphere. American Society for Microbiology; 2018;3. doi:10.1128/MSPHERE.00261-18/ASSET/E79A63FC-67B3-4F2D-8059-10C711481BE6/ASSETS/GRAPHIC/SPH0031825740005.JPEG PMID:29950381

17. Macfarlane S, Macfarlane GT. Regulation of short-chain fatty acid production. Proc Nutr Soc. Proc Nutr Soc; 2003;62: 67–72. doi:10.1079/PNS2002207 PMID:12740060

18. Kondepudi KK, Ambalam P, Nilsson I, Wadström T, Ljungh Å. Prebiotic-non-digestible oligosaccharides preference of probiotic bifidobacteria and antimicrobial activity against Clostridium difficile. Anaerobe. Anaerobe; 2012;18: 489–497. doi:10.1016/J.ANAEROBE.2012.08.005 PMID:22940065

19. McDonald JAK, Mullish BH, Pechlivanis A, Liu Z, Brignardello J, Kao D, et al. Inhibiting Growth of Clostridioides difficile by Restoring Valerate, Produced by the Intestinal Microbiota. Gastroenterology. Gastroenterology; 2018;155: 1495-1507.e15. doi:10.1053/J.GASTRO.2018.07.014 PMID:30025704

20. Bloemen JG, Venema K, van de Poll MC, Olde Damink SW, Buurman WA, Dejong CH. Short chain fatty acids exchange across the gut and liver in humans measured at surgery. Clin Nutr. Clin Nutr; 2009;28: 657–661. doi:10.1016/J.CLNU.2009.05.011 PMID:19523724

21. Morrison DJ, Preston T. Formation of short chain fatty acids by the gut microbiota and their impact on human metabolism. Gut Microbes. Gut Microbes; 2016;7: 189–200. doi:10.1080/19490976.2015.1134082 PMID:26963409

22. Shimotoyodome A, Meguro S, Hase T, Tokimitsu I, Sakata T. Short chain fatty acids but not lactate or succinate stimulate mucus release in the rat colon. Comp Biochem Physiol A Mol Integr Physiol. Comp Biochem Physiol A Mol Integr Physiol; 2000;125: 525–531. doi:10.1016/S1095-6433(00)00183-5 PMID:10840229

23. Hatayama H, Iwashita J, Kuwajima A, Abe T. The short chain fatty acid, butyrate, stimulates MUC2 mucin production in the human colon cancer cell line, LS174T. Biochem Biophys Res Commun. Biochem Biophys Res Commun; 2007;356: 599–603. doi:10.1016/J.BBRC.2007.03.025 PMID:17374366

24. Usami M, Kishimoto K, Ohata A, Miyoshi M, Aoyama M, Fueda Y, et al. Butyrate and trichostatin A attenuate nuclear factor kappaB activation and tumor necrosis factor alpha secretion and increase prostaglandin E2 secretion in human peripheral blood mononuclear cells. Nutr Res. Nutr Res; 2008;28: 321–328. doi:10.1016/J.NUTRES.2008.02.012 PMID:19083427

25. Psichas A, Sleeth ML, Murphy KG, Brooks L, Bewick GA, Hanyaloglu AC, et al. The short chain fatty acid propionate stimulates GLP-1 and PYY secretion via free fatty acid receptor 2 in rodents. Int J Obes (Lond). Int J Obes (Lond); 2015;39: 424–429. doi:10.1038/IJO.2014.153 PMID:25109781

26. Kim MH, Kang SG, Park JH, Yanagisawa M, Kim CH. Short-chain fatty acids activate GPR41 and GPR43 on intestinal epithelial cells to promote inflammatory responses in mice. Gastroenterology. Gastroenterology; 2013;145. doi:10.1053/J.GASTRO.2013.04.056 PMID:23665276

27. Cummings JH, Pomare EW, Branch HWJ, Naylor CPE, MacFarlane GT. Short chain fatty acids in human large intestine, portal, hepatic and venous blood. Gut. Gut; 1987;28: 1221–1227. doi:10.1136/GUT.28.10.1221 PMID:3678950

28. Rolfe RD. Role of volatile fatty acids in colonization resistance to Clostridium difficile. Infect Immun. Infect Immun; 1984;45: 185–191. doi:10.1128/IAI.45.1.185-191.1984 PMID:6735467

29. Theriot CM, Koenigsknecht MJ, Carlson PE, Hatton GE, Nelson AM, Li B, et al. Antibiotic-induced shifts in the mouse gut microbiome and metabolome increase susceptibility to Clostridium difficile infection. Nat Commun. Nat Commun; 2014;5. doi:10.1038/NCOMMS4114 PMID:24445449

30. Ritsema T, Smeekens S. Fructans: beneficial for plants and humans. Curr Opin Plant Biol. Curr Opin Plant Biol; 2003;6: 223–230. doi:10.1016/S1369-5266(03)00034-7 PMID:12753971

31. Nakamura S, Nakashio S, Yamakawa K, Tanabe N, Nishida S. Carbohydrate fermentation by Clostridium difficile. Microbiol Immunol. Microbiol Immunol; 1982;26: 107–111. doi:10.1111/J.1348-0421.1982.TB00159.X PMID:6806571

32. Lammens W, Le Roy K, Schroeven L, Van Laere A, Rabijns A, Van Den Ende W. Structural insights into glycoside hydrolase family 32 and 68 enzymes: functional implications. J Exp Bot. Oxford Academic; 2009;60: 727–740. doi:10.1093/JXB/ERN333 PMID:19129163

33. Louis P, Flint HJ. Formation of propionate and butyrate by the human colonic microbiota. Environ Microbiol. Environ Microbiol; 2017;19: 29–41. doi:10.1111/1462-2920.13589 PMID:27928878

34. Ríos-Covián D, Ruas-Madiedo P, Margolles A, Gueimonde M, De los Reyes-Gavilán CG, Salazar N. Intestinal Short Chain Fatty Acids and their Link with Diet and Human Health. Front Microbiol. Front Microbiol; 2016;7. doi:10.3389/FMICB.2016.00185 PMID:26925050

35. Gregory AL, Pensinger DA, Hryckowian AJ. A short chain fatty acid–centric view of Clostridioides difficile pathogenesis. PLOS Pathog. Public Library of Science; 2021;17: e1009959. doi:10.1371/JOURNAL.PPAT.1009959 PMID:34673840

36. Rossi M, Corradini C, Amaretti A, Nicolini M, Pompei A, Zanoni S, et al. Fermentation of fructooligosaccharides and inulin by bifidobacteria: a comparative study of pure and fecal cultures. Appl Environ Microbiol. Appl Environ Microbiol; 2005;71: 6150–6158. doi:10.1128/AEM.71.10.6150-6158.2005 PMID:16204533

37. Šuligoj T, Vigsnæs LK, Van den Abbeele P, Apostolou A, Karalis K, Savva GM, et al. Effects of Human Milk Oligosaccharides on the Adult Gut Microbiota and Barrier Function. Nutrients. Nutrients; 2020;12: 1–21. doi:10.3390/NU12092808 PMID:32933181

38. Sprenger N, Tytgat HL, Binia A, Austin S, Singhal A. Biology of human milk oligosaccharides: from Basic Science to Clinical Evidence. J Hum Nutr Diet. J Hum Nutr Diet; 2022; doi:10.1111/JHN.12990 PMID:35040200

39. Seekatz AM, Theriot CM, Rao K, Chang YM, Freeman AE, Kao JY, et al. Restoration of short chain fatty acid and bile acid metabolism following fecal microbiota transplantation in patients with recurrent Clostridium difficile infection. Anaerobe. Anaerobe; 2018;53: 64–73. doi:10.1016/J.ANAEROBE.2018.04.001 PMID:29654837

40. Sun Y, O’Riordan MXD. Regulation of bacterial pathogenesis by intestinal short-chain Fatty acids. Adv Appl Microbiol. Adv Appl Microbiol; 2013;85: 93–118. doi:10.1016/B978-0-12-407672-3.00003-4 PMID:23942149

41. Shepherd ES, Deloache WC, Pruss KM, Whitaker WR, Sonnenburg JL. An exclusive metabolic niche enables strain engraftment in the gut microbiota. Nat 2018 5577705. Nature Publishing Group; 2018;557: 434–438. doi:10.1038/s41586-018-0092-4 PMID:29743671

42. Vinolo MAR, Rodrigues HG, Nachbar RT, Curi R. Regulation of inflammation by short chain fatty acids. Nutrients. Nutrients; 2011;3: 858–876. doi:10.3390/NU3100858 PMID:22254083

43. Fachi JL, D. J, Felipe S, Passariello L, Dos A, Farias S, et al. Butyrate Protects Mice from Clostridium difficile-Induced Colitis through an HIF-1-Dependent Mechanism. Cell Rep. 2019;27: 750–761. doi:10.1016/j.celrep.2019.03.054

44. Litvak Y, Byndloss MX, Bäumler AJ. Colonocyte metabolism shapes the gut microbiota. Science (80-). American Association for the Advancement of Science; 2018;362. doi:10.1126/SCIENCE.AAT9076/ASSET/3E223686-1C11-4413-8AA2-6A69D3F5F7E0/ASSETS/GRAPHIC/362_AAT9076_F3.JPEG PMID:30498100

45. Zhang W, Yan J, Wu L, Yu Y, Ye RD, Zhang Y, et al. In vitro immunomodulatory effects of human milk oligosaccharides on murine macrophage RAW264.7 cells. Carbohydr Polym. Carbohydr Polym; 2019;207: 230–238. doi:10.1016/J.CARBPOL.2018.11.039 PMID:30600004

46. Holscher HD, Davis SR, Tappenden KA. Human milk oligosaccharides influence maturation of human intestinal Caco-2Bbe and HT-29 cell lines. J Nutr. J Nutr; 2014;144: 586–591. doi:10.3945/JN.113.189704 PMID:24572036

47. Koenigsknecht MJ, Theriot CM, Bergin IL, Schumacher CA, Schloss PD, Young VB. Dynamics and establishment of Clostridium difficile infection in the murine gastrointestinal tract. Infect Immun. Infect Immun; 2015;83: 934–941. doi:10.1128/IAI.02768-14 PMID:25534943

48. Kumar N, Browne HP, Viciani E, Forster SC, Clare S, Harcourt K, et al. Adaptation of host transmission cycle during Clostridium difficile speciation. Nat Genet 2019 519. Nature Publishing Group; 2019;51: 1315–1320. doi:10.1038/s41588-019-0478-8 PMID:31406348

49. He M, Sebaihia M, Lawley TD, Stabler RA, Dawson LF, Martin MJ, et al. Evolutionary dynamics of Clostridium difficile over short and long time scales. Proc Natl Acad Sci U S A. National Academy of Sciences; 2010;107: 7527–7532. doi:10.1073/PNAS.0914322107/SUPPL_FILE/PNAS.200914322SI.PDF PMID:20368420

50. Sebaihia M, Wren BW, Mullany P, Fairweather NF, Minton N, Stabler R, et al. The multidrug-resistant human pathogen Clostridium difficile has a highly mobile, mosaic genome. Nat Genet. Nat Genet; 2006;38: 779–786. doi:10.1038/NG1830 PMID:16804543

51. Han J, Lin K, Sequeira C, Borchers CH. An isotope-labeled chemical derivatization method for the quantitation of short-chain fatty acids in human feces by liquid chromatography-tandem mass spectrometry. Anal Chim Acta. Anal Chim Acta; 2015;854: 86–94. doi:10.1016/J.ACA.2014.11.015 PMID:25479871

